# Identification, presence, and possible multifunctional regulatory role of invertebrate gonadotropin-releasing hormone/corazonin molecule in the great pond snail (*Lymnaea stagnalis*)

**DOI:** 10.1101/2020.03.01.971697

**Authors:** István Fodor, Zita Zrinyi, Péter Urbán, Róbert Herczeg, Gergely Büki, Joris M. Koene, Pei-San Tsai, Zsolt Pirger

## Abstract

In the last years, the interpretation of gonadotropin-releasing hormone (GnRH) neuropeptide superfamily has changed tremendously. One main driver is the investigation of functions and evolutionary lineage of previously identified molluscan GnRH molecules. Emerging evidence suggests not only reproductive, but also diverse biological effects of these molecules and proposes they should most likely be called corazonin (CRZ). Clearly, a more global understanding necessitates further exploration of species-specific functions and structure of invGnRH/CRZ peptides. Towards this goal, we have identified the full-length cDNA of invGnRH/CRZ peptide in an invertebrate model species, the great pond snail *Lymnaea stagnalis*, termed ly-GnRH/CRZ, and characterized the transcript and peptide distribution in the central nervous system (CNS) and peripheral organs. Our results are consistent with previous data that molluscan GnRHs are more related to CRZs and serve diverse functions. For this, our findings support the notion that peptides originally termed molluscan GnRH are multifunctional modulators and that nomenclature change should be taken into consideration.

## 1. INTRODUCTION

The gonadotropin-releasing hormone (GnRH) is an ancient neuropeptide superfamily whose origin predates the protostome-deuterostome split (Lindemans et al., 2011; Plachetzki et al., 2016; Sakai et al., 2017). Currently, the GnRH superfamily consists of five families: 1) vertebrate GnRH; 2) adipokinetic hormone (AKH); 3) corazonin (CRZ); 4) AKH/CRZ-related peptide (ACP), and 5) invertebrate GnRH/CRZ (invGnRH/CRZ; previously called invGnRH) (Sakai et al., 2017; Tsai, 2018). Through 700 million years of metazoan evolution, these peptides underwent several changes, with only the vertebrate GnRH becoming highly specialized in reproductive activation by stimulating the synthesis and release of luteinizing hormone and follicle-stimulating hormone (Holland and Sower, 2010). For invertebrates, in contrast, AKH, CRZ, ACP and invGnRH/CRZ are shown to be multifunctional modulators of different physiological and behavioral processes (Sakai et al., 2017). Among these peptides, the nomenclature of invGnRH/CRZ molecules has led to some confusion regarding their evolutionary lineage and function (Tsai, 2018). In other words, invGnRH/CRZ molecules, initially named invGnRH (e.g., oct-GnRH, ap-GnRH), may not assume a specialized reproductive role as seen in vertebrate GnRH. Indeed, emerging evidence suggests diverse biological effects of invGnRH/CRZ that include not only reproductive, but also motor, behavioral, and neuromodulatory functions (Iwakoshi-Ukena et al., 2004; Jung et al., 2014; Kavanaugh and Tsai, 2016; Minakata et al., 2009; Nagasawa et al., 2015; Sun and Tsai, 2011; Treen et al., 2012; Tsai et al., 2010). A more global understanding of invGnRH/CRZ evolution necessitates further exploration of species-specific functions and structure of invGnRH/CRZ molecules.

The overarching goal of the present study was to identify and describe the invGnRH/CRZ molecule in a freshwater great pond snail, *Lymnaea stagnalis*, named ly-GnRH/CRZ. For decades, *L. stagnalis* has been a widely used model species for the study of invertebrate neurobiology (Benjamin, 2008; Feng et al., 2009; Kemenes and Benjamin, 2009; Pirger et al., 2014a; Rivi et al., 2020), neurodegeneration processes (Ford et al., 2017; Maasz et al., 2017), aging (Hermann et al., 2007; Pirger et al., 2014b), environmental risk assessment (Amorim et al., 2019; Bouetard et al., 2014; Charles et al., 2016; Pirger et al., 2018; Zrinyi et al., 2017), and the evolutionary consequence of hermaphroditism (reviewed in (Koene, 2017). *L. stagnalis* possesses unique features such as a well-defined central nervous system (CNS), clearly-defined behaviors, and accessible anatomy that can facilitate the characterization of invGnRH/CRZ. To accomplish our goal, we first identified the coding sequence of prepro-ly-GnRH/CRZ. Subsequently, reverse-transcription (RT) polymerase chain reaction (PCR) and immunohistochemistry (IHC) were performed to examine the anatomical distribution and cellular localization of ly-GnRH/CRZ transcript and peptide, respectively. Our data revealed the presence of an invGnRH/CRZ preprohormone in *L. stagnalis.* Further, the mRNA and peptide distribution patterns were consistent with the role of ly-GnRH/CRZ as a neuropeptide with diverse functions.

## 2. MATERIALS AND METHODS

### 2.1. Experimental animals

Adult (5-6 months old) specimens of the pond snail, *L. stagnalis*, originating from our laboratory-bred stocks, were randomly selected for use in experiments. Snails were kept in large holding tanks containing oxygenated low-copper artificial snail water at a constant temperature of 20°C (±1.5 °C) and on light:dark regime of 12 h:12 h. Snails were fed lettuce *ad libitum* three times a week. All procedures on snails were performed according to the protocols approved by the Scientific Committee of Animal Experimentation of the Balaton Limnological Institute (VE-I-001/01890-10/2013). Efforts were made to minimize the number of animals used in the experiments.

### 2.2. *In silico* searches, cDNA sequencing, and bioinformatics analysis

*In silico* searches of *L. stagnalis* CNS transcriptome and genome were conducted with TBLASTN (NCBI). First, the coding sequence of ap-GnRH/CRZ (GnRH/CRZ of *Aplysia californica;* NM_001204553.1) was used as the search query and identified a 272 bp long *L. stagnalis* cDNA fragment in the genome consortium data (referred to as Lsta_scaffold164; genome consortium publication in preparation), indicating the presence of GnRH/CRZ in *L. stagnalis.*

For RNA sample collection, the whole CNS was dissected from the snails (n=10) and homogenized using a TissueLyser LT (QIAGEN) in TRI reagent (#93289, Sigma-Aldrich). RNA was isolated with Direct-zol^TM^ RNA MiniPrep (#R2050, Zymo Research) following manufacturer’s instructions. The RNA was quantified by Qubit BR RNA Kit (#Q10211, ThermoFisher), and the quality was checked on Agilent Bioanalyzer 2100 using RNA 6000 Nano Kit (#5067-1511, Agilent).

Nanopore sequencing was used for identification of ly-GnRH/CRZ whole transcript. The library was prepared using cDNA-PCR Kit (#SQK-PCS108, Oxford Nanopore Technologies) according to the description of manufacturer. The sample was sequenced on a MinION device with R9.4.1 flowcells (#FLO-MIN106). Base calling was performed using Guppy v3.2.2 software. Adapters were trimmed with Porechop v0.2.4 (Wick, 2018), and sequences with internal adapters, indicative of chimera reads, were also split with Porechop. Reads were assembled with CLC Genomics Workbench v12.0.3 software *de novo* pipeline (QIAGEN). Consensus sequence was called and manually corrected also within CLC Genomics Workbench. The identified ly-GnRH/CRZ mRNA sequence was submitted to the GenBank Nucleotide database with the accession number MN385595.

### 2.3. Phylogenetic analysis

The alignment used to generate the maximum likelihood tree consisted of 56 amino acid (AA) sequences of GnRH superfamily preprohormones retrieved from GenBank and the deduced AA sequence of ly-GnRH/CRZ preprohormone (see Supplementary information). The alignment was performed using ClustalW with BLOSUM62 substitution matrix in Molecular Evolutionary Genetics Analysis v7 software (Kumar et al., 2016). Alignment was then analyzed to get the best fitting model, which was determined to be LG with gamma-distributed rates. Bootstrapping support for the tree was conducted with 1000 bootstrap replicates.

### 2.4. RT-PCR analysis of ly-GnRH/CRZ expression in central and peripheral tissues

Total RNA from the CNS (buccal, cerebral, pedal, pleural, parietal, and visceral ganglia) and peripheral organs (heart, and ovotestis) was isolated as presented above. After the RNA preparation, an additional DNase treatment was performed using TURBO DNA-free™ Kit (#AM1907, Thermo Fisher Scientific) following manufacturer’s instructions.

The RevertAid H Minus First Strand cDNA Synthesis Kit (#K1631, Thermo Fisher Scientific) was used for RT, applying random hexamer primers and 3 μL volume from the total RNA sample.

PCR primers were designed for the identified coding region of ly-GnRH/CRZ using SnapGene® Viewer software (GSL Biotech, Chicago, IL, version 4.1.7). The applied primer set was as follows: the forward primer for ly-GnRH: 5’-CCT TGT CCT CCT GGC TGT AGT G - 3’; the reverse primer for ly-GnRH: 5’-GAT GTG GCC GGT GTA CGA TGG - 3’ (Integrated DNA Technologies, Leuven, Belgium). As a control to ensure the quality of RT, actin was also amplified for all examined tissues. The applied primer set for actin was as follows: the forward primer: 5’-TCC CTT GAG AAG AGC TAC GAG C - 3’; and the reverse primer: 5’-GAG TTG TAG GTG GTT TCG TGG - 3’ (Integrated DNA Technologies, Leuven, Belgium).

The PCR reaction was performed in 20 μL reaction volume at 95 °C for 4 min (initial denaturation) followed by 35 cycles of 95 °C for 30 sec (denaturation), 55 °C for 30 sec (annealing) and 72 °C for 10 sec (extension) by a T1 Thermocycler PCR device (Biometra®, Goettingen, Germany). PCR product was checked by agarose gel-electrophoresis using 2% gel (SeaKem LE agarose) and GeneRuler^TM^ 100 bp Plus DNA ladder (#SM0328, Thermo Fisher Scientific).

### 2.5. Immunohistochemistry

CNS, heart, and ovotestis were dissected from individual snails and pinned out on a Sylgard-coated dish containing 4% paraformaldehyde in 0.1 M phosphate-buffered saline (PBS, pH=7.4) for overnight at 4°C. After washing with PBS, fixed tissues were cryoprotected in 20% glucose solution for 4 hours at room temperature and embedded into cryomatrix (#6769006, Thermo Scientific). A series of 12-14 μm-thick cryostat sections were cut, thaw-mounted onto gelatin-aluminium-coated slides, and processed for IHC as follows: 1) washing with PBS containing 0.25% TritonX-100 for 3 x 10 min; 2) incubation with a rabbit anti-ap-GnRH/CRZ antiserum (#AS203-2, EZbiolab; antigen was a synthetic undecapeptide: CNYHFSNGWYA-amide) (Tsai et al., 2010) diluted 1:750 for 24 hours at 4°C; 3) PBS washing for 2 x15 min; 4) incubation with a donkey anti-rabbit IgG conjugated with NorthernLights^™^ NL557 (#NL004, R&D System) diluted 1:1000 for 2 hours at 4°C. After washing with modified Dulbeco’s PBS (DPBS), the nuclei were stained with Hoechst 33342 (1 μg/mL in DPBS; #62249, Thermo Scientific) for 10 min at room temperature. The slides were washed with DPBS and cover-slipped with fluorescent mounting medium (#S3023, Dako). The stained tissues were analyzed with a TCS SP8 DMI laser confocal scanning microscope (Leica Microsystems, Germany) equipped with appropriate wavelength-filter configuration settings and a transmitted light detector (BioMarker Ltd, Hungary). The necessary number of optical line sections (15-30) with 0.5-0.8 μm step-size were made to capture of all visualized details. Image processing was performed by LasX software (Leica Microsystems, Germany). Because the primary antiserum was generated against ap-GnRH/CRZ conjugated with hemocyanin (KLH) from giant keyhole limpet (*Megathura crenulata*), to abolish cross-reaction to hemocyanin in tissues of *L. stagnalis* the antiserum was preadsorbed overnight against KLH (#H7017, Sigma). In some cases, the antibody was preadsorbed with 10 μM of synthetic ly-GnRH/CRZ (pQNYHFSNGWYA-amide) overnight at 4°C and applied to the sections.

## 3. RESULTS

### 3.1. Nucleotide sequence of ly-GnRH/CRZ preprohormone

Our *in silico* search in the *L. stagnalis* sequence data revealed a 272-bp positive hit (referred to as Lsta_scaffold164) containing a sequence of 32 nucleotides with a 100% match to the query, thus identifying the potential GnRH/CRZ coding sequence. We termed this sequence ly-GnRH/CRZ. Using nucleotide sequencing, we determined the complete coding sequence. The full-length ly-GnRH/CRZ preprohormone spans 1225 bp (Fig. 1). Based on the data from scientific literature (Iwakoshi et al., 2002; Tsai and Zhang, 2008; Zhang et al., 2008), we identified the signal peptide, the mature peptide with the signature tribasic cleavage site and an alpha amidation signal (QNYHFSNGWYAGKKR), and the ly-GnRH/CRZ-associated peptide (Fig. 1). As expected, the mature peptide sequence is highly conserved, while the signal peptide and GnRH/CRZ-associated peptide show high variability.

**Figure 1 –.**
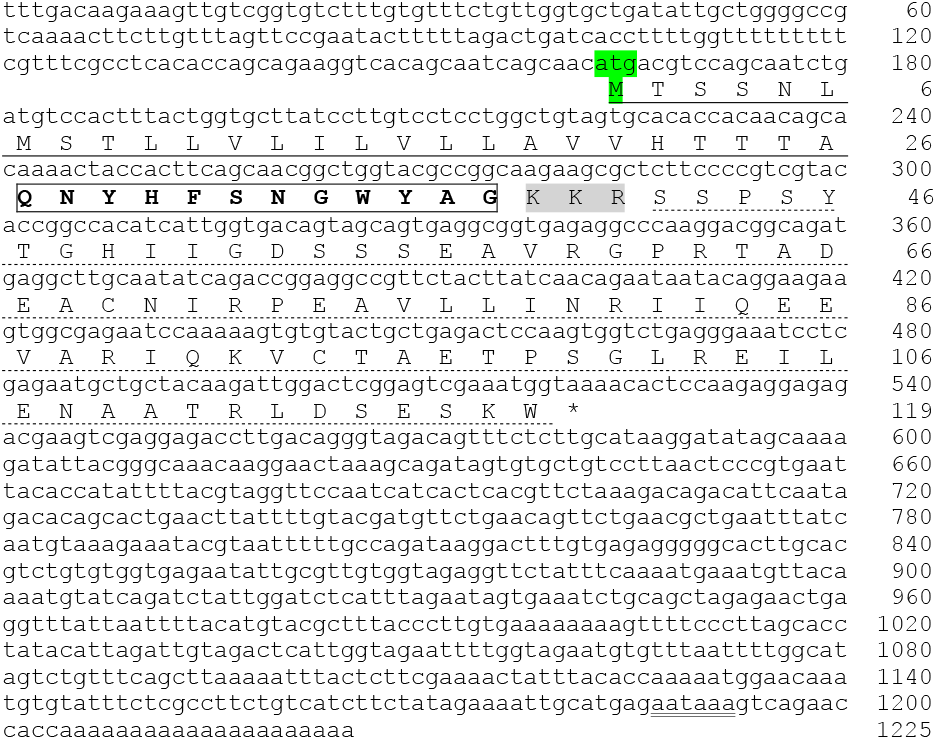
Nucleotide and deduced AA sequences of ly-GnRH/CRZ preprohormone (Genbank Accession #MN385595). The nucleotides (upper row) and AAs (lower row) are numbered accordingly. The putative signal peptide region is underlined (solid). The putative mature ly-GnRH/CRZ peptide with an alpha amidation signal is boxed, and the KKR tribasic cleavage site is shaded. The putative ly-GnRH/CRZ-associated peptide is underlined (dashed). The asterisk (*) denotes the stop codon. The nucleotides corresponding to the polyadenylation signal (ATTAAA) are double-underlined.

### 3.2. Phylogenetic analysis

To update the established evolutionary relationships of the GnRH superfamily with newly annotated sequences including ly-GnRH/CRZ, we performed a phylogenetic analysis. The results of this are presented in Fig. 2, and the segregated clusters from top to bottom are as follows:

1. Two clusters of AKHs from mollusks, arthropods, annelids, and nematodes. Based on data from the literature, all of molluscan, arthropod and nematode AKHs are true AKH molecules. Five manual corrections are required: *Crassostrea gigas* AKH belongs to molluscan AKH group, *Bombyx mori* ACP to the ACP group, *Branchiostoma floridae* GnRH to the GnRH group, while *Capitella teleta* and *Helobdella robusta* (annelids) AKHs are not true AKH molecules but referred to as AKH/GnRHs. Furthermore, one worm peptide, *Priapulus caudatus* AKH, should belong to AKH cluster.
2. An arthropod ACP cluster. Two manual corrections should also be performed here; *Brachionus calyciflorus* AKH has the consensus sequence for arthropod AKH but not ACP cluster, and *Cionia intestinalis* GnRH belongs to GnRH cluster.
3. An arthropod CRZ cluster.
4. A GnRH/CRZ cluster, including the newly determined ly-GnRH/CRZ (arrowhead) and previously identified molluscan GnRH sequences, possesses both GnRH and CRZ properties.
5. A vertebrate GnRH cluster containing all used sequences from chordates and echinoderms.

**Figure 2 –.**
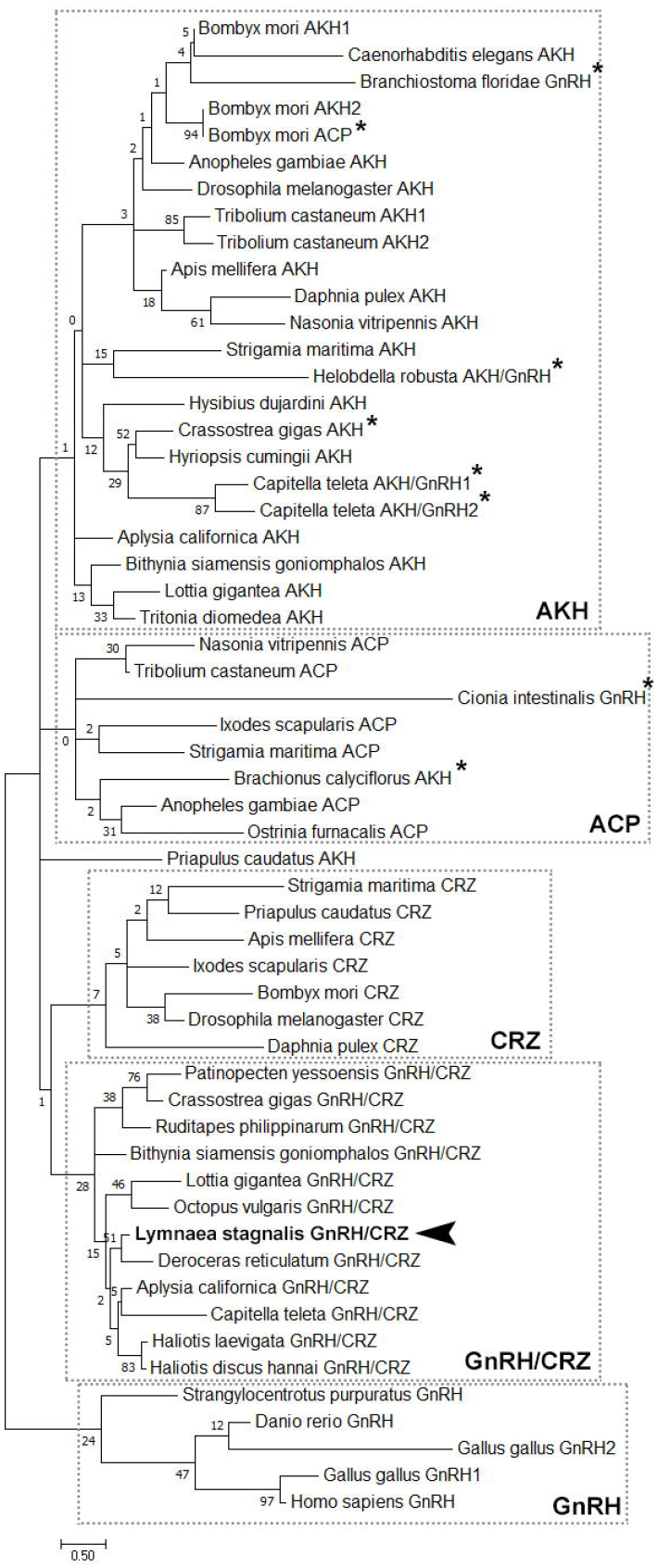
Phylogenetic analysis of CRZ, GnRH, GnRH/CRZ, AKH, and ACP preprohormone sequences derived from protostomes including mollusks, annelids, nematodes and arthropods, and deuterostomes including echinoderms and chordates. The bootstrap values (%) from 1000 replicas are indicated at each branch point, the analysis involved 56 sequences. Sequences indicated by an asterisk belong to a neuropeptide group, which is different from the ones in which they cluster in the analysis.

### 3.3. Localization of ly-GnRH/CRZ transcript in central and peripheral tissues

RT-PCR revealed wide distribution of ly-GnRH/CRZ transcript (Fig. 3). The ly-GnRH/CRZ transcript (109 bp) can be detected in all CNS ganglia (CG, BG, PeG, PlG, PaG, VG) and peripheral organs examined, including the heart (H) and ovotestis (OT) (Fig. 3, top panel). Actin (a housekeeping gene) amplification of RNA samples without RT was used as a negative control to test for genomic DNA contamination (Fig. 3, middle panel). The lower panel amplified actin (140 bp) to ensure RNA and RT quality. Negative controls including no RT (middle panel) and no template (NTC) controls did not yield positive signal.

**Figure 3 –.**
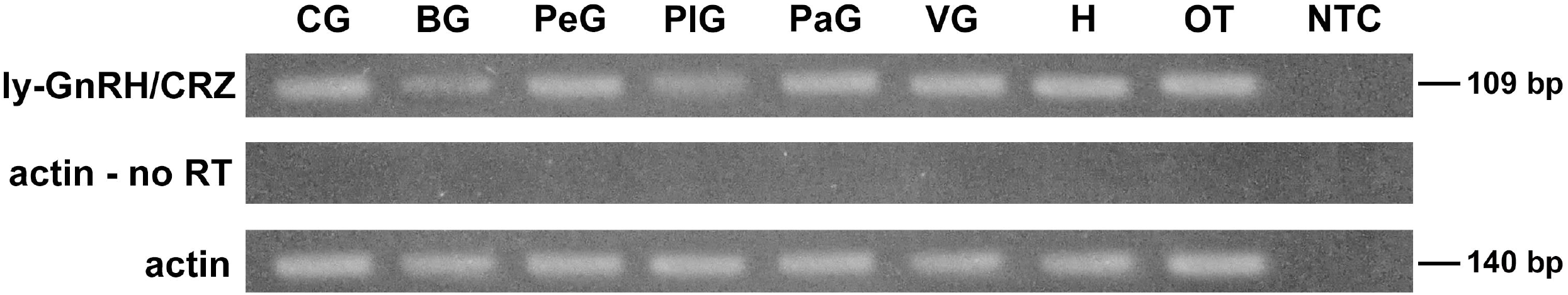
PCR of cDNA synthesized from the CNS (buccal, cerebral, pedal, pleural, parietal, and visceral ganglia) and peripheral tissues (heart, ovotestis). Top panel: PCR of ly-GnRH/CRZ. Middle panel: PCR of ly-actin using RNA samples that have not been reverse transcribed to check for genomic DNA contamination. Bottom panel: PCR of ly-actin to check the quality of RNA samples. The sizes of PCR products are indicated on the right. CG: cerebral ganglia; BG: buccal ganglia; PeG: pedal ganglia; PlG: pleural ganglia; PaG: parietal ganglia; VG: visceral ganglion; H: heart; OT: ovotestis; NTC: no template negative control.

### 3.4. Localization of ly-GnRH/CRZ peptide in central and peripheral tissues

IHC revealed the presence of ly-GnRH/CRZ peptide in all CNS ganglia (Fig. 4A). Large-(50-80 μm), medium-(30-50 μm) and small-sized (10-30 μm) ly-GnRH/CRZ-immunoreactive (ir) neurons were observed. The distribution of ly-GnRH/CRZ-ir neurons was highly variable in different ganglia (Fig. 4A). In the pleural (PlG) and parietal (PaG) ganglia, only a few ly-GnRH/CRZ-ir neurons were detected, but labeled neurons were more abundant in the buccal (BG), cerebral (CG), and pedal ganglia (PeG) (Fig. 4A). The BG contained several small-and medium-sized ly-GnRH/CRZ-ir neurons largely concentrated around the unlabeled B1 and B2, furthermore, around the immunopositive large-sized B3 and B4 feeding motoneurons (Fig. 4A and B). In the CG, small-sized ly-GnRH/CRZ-ir neurons were detected in the metacerebrum located close to the ventral lobe (vl), in the anterior lobe (al), and a group of medium-sized ly-GnRH/CRZ-ir neurons were observed in the A-cluster (Acl) (Fig. 4A and C). Furthermore, caudo-dorsal cells (CDCs), the reproductive neuroendocrine cells, also displayed immunoreactivity (Fig. 4A and C). The PeG contained several small-and medium-sized immunopositive neurons in the A-(Acl) and E (Ecl)-clusters. The left serotonergic giant cell (LPeD1) and the functionally identified right dopaminergic giant neuron (RPeD1) were also immunopositive (Fig 4A and D). Lastly, some medium-and large-sized ly-GnRH/CRZ-ir neurons were observed in the J cells and M-cluster (Mcl) of the visceral ganglion (VG), and in the B-cluster (Bcl) of the PaG (Fig. 4A and E). Generally, a strong presence of ly-GnRH/CRZ-immunoreactivity was detected in the neuropil regions throughout the CNS (Fig. 4C-E).

**Figure 4 –.**
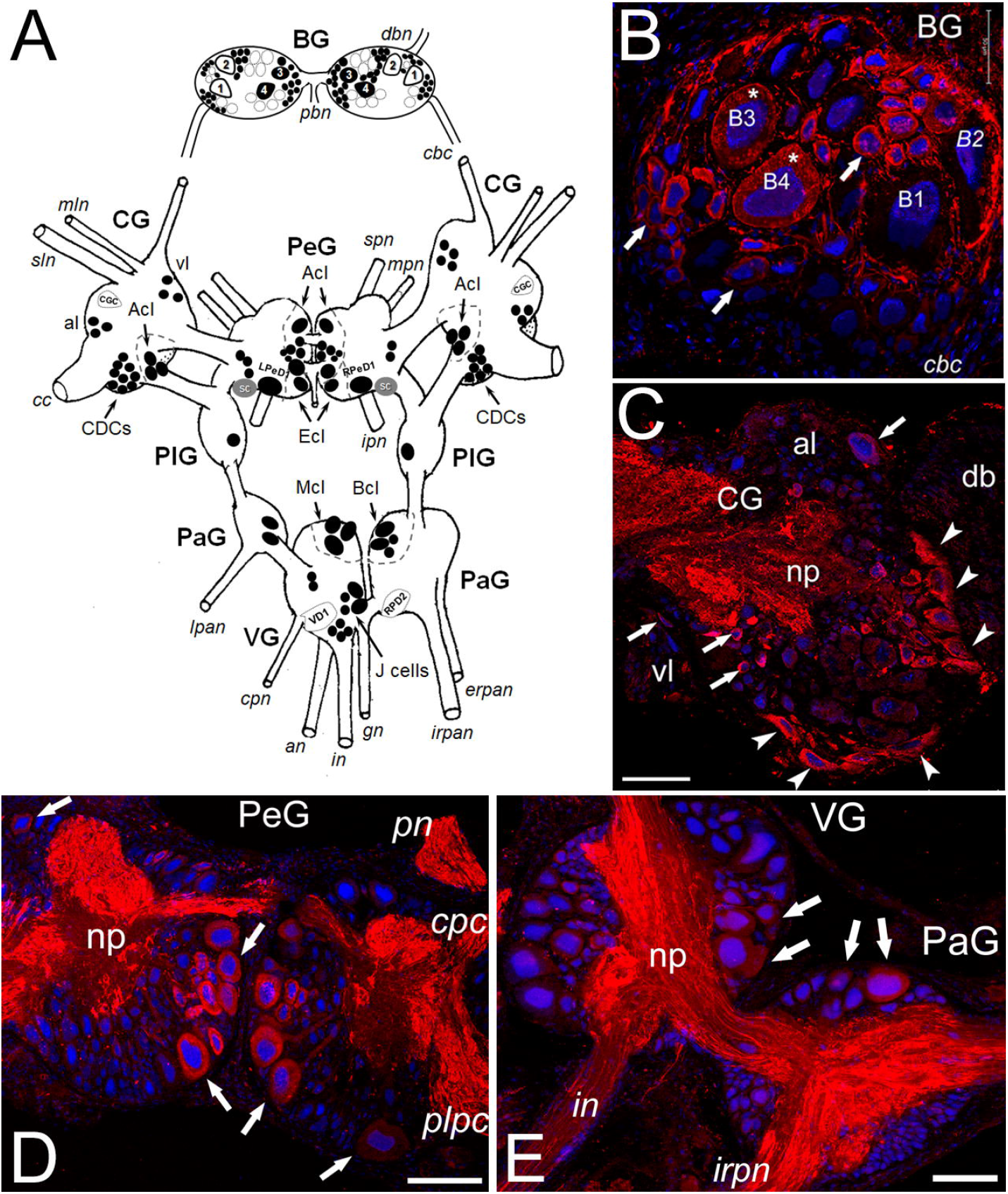
Schematic drawing and representative micrographs showing ly-GnRH/CRZ-ir neuronal elements in different ganglia of CNS. (A) Schematic map (dorsal view) of the distribution of ly-GnRH/CRZ-ir cells in the buccal (BG), cerebral (CG), pedal (PeG), pleural (PlG), parietal (PaG), and visceral (VG) ganglia of *L. stagnalis.* ly-GnRH/CRZ-ir neurons are indicated with black symbols, while outlined areas (black dashed lines) indicate unlabeled but identified (e.g., CGC, VD1) large neurons providing reference points for anatomical orientation. (B) In the right BG small-and medium-sized unidentified ly-GnRH/CRZ-ir cells (arrows), and large identified immunopositive neurons (stars, B3, B4) can be observed. (C) In the left CG several identified (CDCs, arrowheads) and unidentified (arrows) cell bodies show positive immunosignal. (D, E) Several immunopositive small (10-30 μm), middle (30-50 μm), and large (50-80 μm) neurons (arrows) can be observed in the PeG (A-and E-clusters), VG (M-cluster), and right PaG (B-cluster). Furthermore, intensive immunostaining in the neuropiles, connectives and peripheral nerves can be observed. Abbreviations: dbn, dorsobuccal nerve; pbn, postbuccal nerve; cbc, cerebro-buccal connective; mln, median lip nerve; sln, superior lip nerve; vl, ventral lobe; al, anterior lobe; cc, cerebral commissure; CDCs, caudo-dorsal cells; spn, superior pedal nerve; mpn, median pedal nerve; ipn, inferior pedal nerve; Acl, A-cluster; Ecl, E-cluster; SC, statocyst; lpan, left parietal nerve, cpn, cutaneous pillar nerve; an, anal nerve; in, intestinal nerve; gn, genital nerve; irpan, internal right parietal nerve; erpan, external right parietal nerve; Mcl, M-cluster; Bcl, B-cluster; B1-B4, identified buccal feeding motoneurons; db, dorsal body; np, neuropile; pn, pedal nerve; cpc, cerebro-pedal connective; plpc, pleuro-pedal connective. Scale bars=100 μm.

Among peripheral tissues, we have investigated the heart and the ovotestis. Confocal projections showed abundant ly-GnRH/CRZ-ir fibers on the surface of the heart atria running perpendicularly to the longitudinal axis of the muscle fibers; however, the heart muscle fibers themselves were not immunopositive (Fig. 5A1 and A2). In ovotestis, immunoreactivity was not observed (Fig. 5B). The immunostaining was completely abolished when the antiserum was preadsorbed with 10 μM ly-GnRH/CRZ (not shown).

**Figure 5 –.**
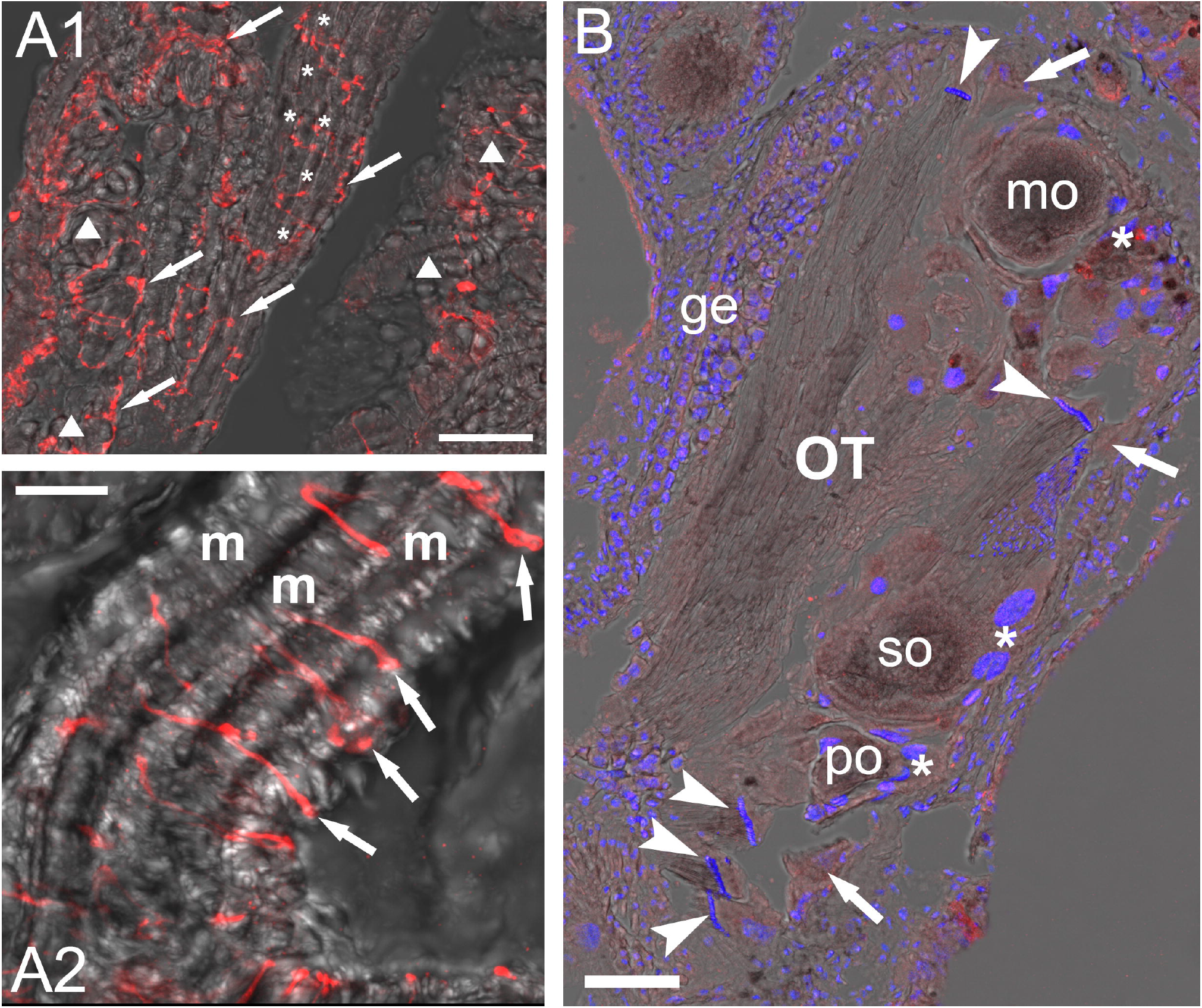
Representative ly-GnRH/CRZ IHC of heart atria (A1-A2) and ovotestis (B). Rich ly-GnRH/CRZ-ir fiber bundles (arrows) can be seen on the surface of transversal (asterisks) and cross-sectional (triangles) heart atria muscles (A1). A2 represents a higher magnification detail of the atria muscle. Immunolabeled axon processes (arrows) running perpendicular the longer, longitudinal axis of the muscle fibers (m). Scale bar=25 μm on A1 and 10 μm on A2. (B) The ovotestis (OT) is devoid of positive signal. Abbreviations: ge, germ epithelium; po, primary oocyte; so, secondary oocyte; mo, mature oocyte; asterisks, follicular cells; arrows, Sertoli cells; arrowheads, spermatozoon.

## 4. DISCUSSION

Previous studies suggested that invGnRH/CRZ molecules, initially named invGnRH, may not assume a specialized reproductive function (Jung et al., 2014; Minakata et al., 2009; Tsai et al., 2010) and may need to be reclassified as CRZ (Hauser and Grimmelikhuijzen, 2014; Plachetzki et al., 2016; Tsai, 2018). The main goal of the present study was to further test this notion by identifying and investigating the invGnRH/CRZ molecule in *L. stagnalis.* The presence of a GnRH-like molecule in *L. stagnalis* has previously been hypothesized (Koene, 2010), but remained unconfirmed until the current report. The identified ly-GnRH/CRZ preprohormone sequence contains the mature peptide with the signature tribasic cleavage site and an alpha amidation signal (QNYHFSNGWYAGKKR). However, the predicted mature peptide sequence awaits validation by mass spectrometry. Based on the literature (Nagasawa et al., 2015), we hypothesize that two forms of ly-GnRH/CRZ may be derived from a single prohormone sequence: a mature amidated undecapeptide (pQNYHFSNGWYA-amide) and a non-amidated precursor dodecapeptide (pQNYHFSNGWYAG-OH).

Our phylogenetic analysis presents two challenges commonly seen in peptide hormone analysis. Firstly, the relatively short GnRH superfamily prepropeptide sequences (58–219 AA) contribute to the lower bootstrap values. Secondly, the conserved mature peptide sequence needed for biological activity (e.g., receptor binding) can cause incorrect clustering (Hauser and Grimmelikhuijzen, 2014). Nevertheless, the phylogenetic tree correlated well with previous studies (Hauser and Grimmelikhuijzen, 2014; Plachetzki et al., 2016; Tsai, 2018). The GnRH/CRZ branching suggests sequences originally termed molluscan GnRHs are no more related to vertebrate GnRH than to AKH, ACP or CRZ.

This is the first study to demonstrate the occurrence and distribution of a snail GnRH/CRZ in the CNS and periphery. Our RT-PCR results show that ly-GnRH/CRZ is ubiquitously present in all central and peripheral tissues examined. However, IHC data show that the simultaneous presence of ly-GnRH/CRZ transcript and peptide occurs predominantly in the CNS. For example, although the ly-GnRH/CRZ transcript can be found in the heart and ovotestis (Fig. 3), the peptide was not observed within the heart muscle proper and absent in the ovotestis (Fig. 5). Similar observations were made in *A. californica* (Jung et al., 2014; Tsai et al., 2010), but the reason for the transcript/peptide mismatch remains to be determined. Proposed explanations include peripheral peptide levels below IHC detection, rapid peripheral peptide turnover or transport from the source tissue, and alternative post-translational processes in the periphery (Jung et al., 2014).

In contrast to *A. californica* in which ap-GnRH/CRZ peptide was found only in pedal and cerebral ganglia, ly-GnRH/CRZ peptide was detected by IHC in all central ganglia of *L. stagnalis*, including functionally and/or anatomically identified neurons (Benjamin, 2008). As such, the possible functions of ly-GnRH/CRZ can be inferred from the distribution and production sites. The B3 and B4 motoneurons in BG (Fig. 4A and B), involved in the execution of feeding (Brierley et al., 1997), showed ly-GnRH/CRZ-ir, suggesting a possible role of ly-GnRH/CRZ in this activity. Furthermore, CV cells in vl of CG (Fig. 4A and C) are well-known for feeding control via the lip motoneurons (Mccrohan, 1984), suggesting a role of this neuropeptide in the motor control of feeding. Interestingly, egg-laying regulatory CDCs of cerebral ganglia also showed ly-GnRH/CRZ-ir (Fig. 4A and C). This contrasts with findings that bag cells of *A. californica*, which are homologous to CDCs and also responsible for egg laying (Conn and Kaczmarek, 1989), are negative for ap-GnRH/CRZ (Zhang et al., 2008). In the PeG, ly-GnRH/CRZ-containing neurons overlap with that of the pedal motor serotonergic neurons (A cluster, Fig. 4A and D) involved in the control of foot cilia (McKenzie et al., 1998), thus predicting a role of ly-GnRH/CRZ in locomotion. The suggested roles of ly-GnRH/CRZ in feeding and locomotion are consistent with previous findings showing ap-GnRH/CRZ modulates these activities in *A. californica* (Jung et al., 2014; Tsai et al., 2010). Furthermore, RPeD1, known to be responsible for respiration and heart control (Benjamin, 2008), is immunopositive. Since ly-GnRH/CRZ-ir J-cells and M-cluster of the VG are also involved in heart control and respiration, the peptide may also play a role in these processes. Supporting a cardio-respiratory role of ly-GnRH/CRZ, ly-GnRH/CRZ-ir fibers are present on the surface of heart atria muscles (Fig. 5A1 and A2). In this respect, it is interesting to note that CRZ molecule was first discovered and named for its ability to stimulate the heart of the American cockroach *(Periplaneta americana)* (Veenstra, 1989). This effect was also documented by the investigation of CRZ of the blood-sucking bug, *Rhodnius prolixus* (Patel et al., 2014). Furthermore, the first described invGnRH/CRZ molecule, the oct-GnRH/CRZ, was also identified in the course of bioassay screenings of peptide fractions with cardio-acceleratory effects (Iwakoshi et al., 2002).

Additional evidence supporting a multifunctional modulatory role of ly-GnRH/CRZ is the observation that some of ly-GnRH/CRZ-ir neurons overlaps with identified serotonergic (e.g., CV cells, A cluster) and dopaminergic (e.g., RPeD1) neurons responsible for feeding, locomotion, heart control, and respiration (Benjamin and Winlow, 1981; Benjamin, 2008; McKenzie et al., 1998; Syed and Winlow, 1991).

In summary, we conclude that ly-GnRH/CRZ, just like ap-GnRH/CRZ, is a multifunctional neuropeptide involved in diverse central functions including feeding, locomotion, heart control, and reproduction. These finding can be confirmed in the future with physiological studies using purified or synthetic ly-GnRH/CRZ peptide. Our results further suggest the term “invGnRH” (e.g., oct-GnRH, ap-GnRH) should be revised to avoid the connotation that these peptides serve specialized reproductive function similar to vertebrate GnRHs (Iwakoshi-Ukena et al., 2004; Jung et al., 2014; Kavanaugh and Tsai, 2016; Minakata et al., 2009; Nagasawa et al., 2015; Sun and Tsai, 2011; Treen et al., 2012; Tsai et al., 2010). Overall, our study reports the presence and peptide distribution of a second invGnRH/CRZ in a gastropod mollusk and adds to a growing body of literature supporting a multifunctional role of invertebrate GnRH/CRZ peptides.

## Funding

This work was supported by National Brain Project (No. 2017-1.2.1-NKP-2017-00002), Bolyai Foundation (No. BO/00952/16/8), and National Science Foundation (IOS-1352944).

## Acknowledgement

Bioinformatic infrastructure was supported by ELIXIR Hungary.

