## Supplementary material for "Identification, presence, and possible multifunctional regulatory role of invertebrate gonadotropin-releasing hormone/corazonin molecule in the great pond snail (*Lymnaea stagnalis*)": Suplementary information: Supplement.docx

**Figure 1.** Amino acid sequences and GenBank accession numbers of the prepropeptides of GnRH superfamily used in the phylogenetic tree analysis. The putative signal peptides (predicted by SignalP) are underlined; the putative active neuropeptides are marked in yellow; glycines used for amidation are shown in blue; green indicates basic cleavage sites flanking the putative neuropeptides.

>Anopheles_gambiae_AKH (XP_563757)

MDTVKLFTVLLICASLMLITEAQLTFTPAWGKRSQGAMGINPLGSTFGQDACKTPVDSLLVIYRMIQAEAQKIVDCSQK

>Apis_mellifera_AKH (AEW68342.1)

MHRKLRPSFFAFSIILLLCLILNTGVEAQLNFSTGWGKRSQRIGVVEWGVRTECATQAKPSVEQLLSVYHLIQIEARKMLDCRKLNE

>Bombyx_mori_AKH1 (NP_001104825.1)

MYKFTILFLVLACFIMAEAQLTFTSSWGGKRAAIAGTVSCRNDESLASIYKLIQNEAEKLLLCQKP

>Bombyx_mori_AKH2 (NP_001124365.1)

MGRALVLVLILSAALLVCEAQLTFTPGWGQGKRSEATDYRNDGCSSEDSVYTIYKLIKNEAEKFLACRS

>Daphnia_pulex_AKH (EFX68649.1)

MANHRILILTLLMIGLASAQVNFSTSWGKRSPSTSTKAAEPPSAPSYRQNFHSKKVEPGTLETLPNNQHLPESFDTVSSTIYDDAEEQRISISLPSPCLSILKSLLLVNQIVEFKNSPLDGRMHRFKIENLFPLPNRTCRLYIRR

>Drosophila_melanogaster_AKH (P61855)

MNPKSEVLIAAVLFMLLACVQCQLTFSPDWGKRSVGGAGPGTFFETQQGNCKTSNEMLLEIFRFVQSQAQLFLDCKHRE

>Nasonia_vitripennis_AKH (ADM26613)

MNCRSLLALGLCLCVVLLQTRAAEGQLNFSTGWGKRSSHLLQPARASSSSSSSSSSSSTSAAAAAAASSGIGRPRQQADLDYFLQRYYRRLRKIEAQRLANSQV

>Strigamia_maritima_AKH (from WGS, AFFK01019834.1)

MTKFTWLSMTLLVLMVFITVDVNGQINFSPGWGQGKRSLSDDKPVNGYSDCSETMIEVYRLLK

>Tribolium_castaneum_AKH1 (ABN79648.1)

MSRMFLIVVLIAFVGVCTAQLNFSTDWGKRSGSSAGSDANNCKEPVETIMLIYKIIQVSFFNIPNSPQQIFAE

>Tribolium_castaneum_AKH2 (ABN79648.1)

MHRVLLTVLLITIVGLCAAQLNFTPNWGKRAPEGESNRCKESVDTIMLIYKIIQVFANYRNFFSHAIFLPERGAEISGL

>Hysibius_dujardini_AKH (from EST, CK326138.1)

MGSAVQSLGLGFLGALLLVQVLFFQVPTANSQLSFSTGWGHGKRSGNVPVNLAPADYRNGKANGDLCSAAYNEMAVARIQELIQEEGFRQLACRKR

>Caenorhabditis_elegans_AKH (NP_500770.1)

MQLYVVLCFLVLLGLSAGQMTFTDQWTKKRATLHKQLPVVTPEEPICPSDRVQAVFEQLDQLQKAQQRLTEYLASCAYPVEVPQKAEKM

>Aplysia_calofirnica_AKH (NP_001268793)

MESSSILLILVVLVIGTSTCLAQIHFSPDWGTGKRAVSTVTEKEIPHCWQIADKEIIDIMLLIQRTAKKLSSCLNTCPEL

>Bithynia_siamensis_goniomphalos_AKH (from EST, GAQQ01002646.1)

MAQTTMTSLLAVLLLLSLVLTPVTSQIHFTPGWGSGKRSGRDDPSSSSSSSASSSSHSSSSIPLPFSSIDSCWAEADFRLFLQLARLVKREAKRFSQCLETENSMAENQENGPDFL

>Crassostrea_gigas_AKH (from WGS, AFTI01019272.1)

MLCKFCILAVVAVSLLSLTLGQVSFSTNWGSGKRSSSSFIAPQADDACWTKSNAKLLYDLMQVIQKQVDRLVACQNSNDDLHRIW

>Hyriopsis_cumingii_AKH (from EST, CW693635.1)

MQTTCIIAVTLVALWTFTSAQISFSTNWGSGKRSGYQDLQSAACLDDLEINVMKDLAALIKMEANRAAKCIIDAKTRKLTGVYETPDK

>Lottia_gigantea_AKH (from EST, FC743844.1)

MSLSRNLSVLVCLCCLLSMCLAQIHFSPTWGSGKRSAPVQTYPDSSNTNCYDKLNTKILYQLVKIIKKESEELTACLLQDDDFLRR

>Tritonia_diomedea_AKH (from EST, EV287376.1)

MKIPDMHRTLHCSIPVLVLLMCVCSSLAQIHFSPGWEPGKRSMEEPTRANKMACYDQLDMSLLLDILKIIKRQAEKLSYCTKSCPQL

>Priapulus_caudatus_AKH (from EST, AXZU01076239)

MHCILILATCLLGLYIHESSAQIFFSKGWRGGKRQEAPLMRHPTDVEEDTNQLVTTALGKESLKLGPSTEPLVDVSLRDAKAWHLTYRTPS

>Brachionus_calyciflorus_AKH (from EST, GACQ01031992)

MKNSASVAIICLSVLVVSIVFDKAMCNPQLTFSSDWSGGKRELENSKTISNDKLDLIGKINFH

>Capitella_teleta_AKH/GnRH1 (ELU16520.1)

MKHFTLLLVACIVVAYHMNTASAQFSFSLPGKWGNGKRAFSFSLPGRWGAQGKRASWTGATTDCSRMESDGMMSVYAAIQEEAIKMLECMGKTSADKQENSLVGAPNINATLVLVSINFQKPISMKSDYETSFE

>Capitella_teleta_AKH/GnRH2 (from WGS, AMQN01001996.1)

MNHMMVVLVACSVLAFQLHSAEAQGFSFSLPGKWGGAGKRSGGVPWLKDGLGECGTYDPSLVVELYKIIQAEAFRLNECLMKAQREGN

>Helobdella_robusta_AKH/GnRH (from WGS, AMQN01001996.1)

MNCLALFLACLVLSTVIILPAKSQFSFTPPGKWPFGTGKRDKIISDDDLTSFQEECFKQDPRILSQTFLRLQVQNLKCKF

>Anopheles_gambiae_ACP (XP_563757.1)

MNSISSSRHLAAKLFLLVALCAVLLPVPSAGQVTFSRDWNAGKRAMPDSPVSGVAECSAIWRSVNNLCAAVTKNIQHLTLCETRSLLKSLQTDESSMESNSGNNLPMFSNNHI

>Bombyx_mori_ACP (NP_001124365.1)

MGRALVLVLILSAALLVCEAQLTFTPGWGQGKRSEATDYRNDGCSSEDSVYTIYKLIKNEAEKFLACRS

>Ixodes_scapularis_ACP (from WGS, ABJB010491828.1)

MASVSRAALAFLLLGLLVVQHVQCQITFSKNWQPGKRGEMCSQREAQAVIKLRQVLFR>Nasonia_vitripennis_ACP (NP_001161199.1)

MGRRLSIGLAAAAILFSCMLHFALAQVTFSKGWGPGKRSALYETDCSRVNYKSLALLFHTLLAEVKHLMACDHQATVNYLQSVERQ

>Ostrinia_furnacalis_ACP (from EST, GAQJ01047301.1)

MNTVRCRGVTVVLVCALAMALVSAQITFSRDWTGGKRAAPMAIDCGQFTRLCRHFVHELKQ

>Strigamia_maritima_ACP (from WGS, AFFK01019675.1)

MKWIAVYLLLTIIVLTIVAPVEGQVTFSRDWTPAGKRGMDCGFVKTKLLRDIAVLLQVKYHFAELCMLTLAWLMLDGGQFTEILWTSCDGAFP

>Tribolium_castaneum_ACP (NP_001159497)

MALKFRVFALVAVLVLMAWMFTGTQAQVTFSRDWNPGKRTENTDLHNTLKTASAVCHLLMNQVRQLASCDNNNELEPGATIFSGRR

>Apis_mellifera_CRZ (NP_001012981.1)

MVNSQILILFILSLTITIVMCQTFTYSHGWTNGKRSTSLEELANRNAIQSDNVFANCELQKLRLLLQGNINNQLFQTPCELLNFPKRSFSENMINDHRQPAPTNNNY

>Bombyx_mori_CRZ (BAC66443.1)

MVTNITLILTLMTLASVTAQTFQYSRGWTNGKRDGHKRDELRDEVLERILTPCQLDKLKYVLEGKPLNDRLFVPCDYIEEEVNQPKRYKGERNHELFDVFQ

>Daphnia_pulex_CRZ (ACJ05606.1)

MFINQYVRYSSSIAMAVRLYFVLLLVVVSAMAQTFQYSRGWTNGRKRSDPSFVQQQQWIQRNGHPIVVPAEFRSNSFEDWSRYRINSEKVFLIVCSCVTFSKRDFMLISVGCHDDNR

>Drosophila_melanogaster_CRZ (AAF55046.1)

MLRLLLLPLFLFTLSMCMGQTFQYSRGWTNGKRSFNAASPLLANGHLHRASELGLTDLYDLQDWSSDRRLERCLSQLQRSLIARNCVPGSDFNANRVDPDPENSAHPRLSNSNGENVLYSSANIPNRHRQSNELLEELSAAGGASAEPNVFGKH

>Ixodes_scapularis_CRZ (from WGS, ABJB010772673.1)

MSHYLGLSAVLLVCLAVTAYSQTFQYSRGWTNGKRRVAEMPLVGVPLRASNDHRALDEVLSKFTPRDRIVLERLGHMVRVLDHAEEEQQEY

>Strigamia_maritima_CRZ (from WGS, AFFK01019339.1)

MGFQKTKLLLIVASILVFIICTSGQTFQYSKGWEPGRKRAVDRSYQVRDWDAGRKRENIRGLDSSTAWILGLKRAAFN

>Priapulus_caudatus_CRZ

MSLSRTLTTMLLLAIAALMLSHVTTAQTFHYSRGWEGRKRSDDSMVSSQQRNTKTPLMSILRHRLLKVRYSHSHLTLLDSYFTRPVMP

>Aplysia_californica_GnRH/CRZ (NP_001191482)

MACRITSATTTLFSILLLIVIAELCSAQNYHFSNGWYAGKKRSSASLLRPEALGSSSSSSSNSGLDVTDADSSRGGVSSLSGRGFGVGELRLVDSPCSIRLETLALINKLMQEEAARIQRTCVANVPSGLRELLEGAASKLESENKW

>Lymnaea_stagnalis_GnRZ/CRZ (this paper; MN385595)

MTSSNLMSTLLVLILVLLAVVHTTTAQNYHFSNGWYAGKKRSSPSYTGHIIGDSSSEAVRGPRTADEACNIRPEAVLLINRIIQEEVARIQKVCTAETPSGLREILENAATRLDSESKW

EITRIITTCTNTVNDIADLQ

>Deroceras_reticulatum_GnRZ/CRZ (ARS01378.1)

MVCSKVTYATLAVTILVLALVQHSLSQNYHFSNGWYPGKKRSAATTSAANSDSGRRRGASASGALDEACGISPYALLLINRIVQDEVARLQKSCSPDLPSGFTEILENAANTLESDSKW>Haliotis_laevigata_GnRH/CRZ (AKR13997.1)

MSVLSSQGVTVSVLLLLLTVHAVAGQNYHFSNGWHAGRKRGGDSTSCVFRKDILMLVNKLIMEESSRVAHRCQNAVPHMDFTEPDTVDAESDGQLTDKRWK

>Haliotis_discus hannai_GnRH/CRZ (AZL93822.1)

MSVLSRQGVTVSVLVLAELCSAQNYHFSNGWYAGKKRSCVFRKDILMLVNKLIMEESSRVAHRCQNGVPHMDFTEPDTVEADSARRQETWRGQHQLRLQERHPHACQSSASLLRPEALGSSSSSSSNSGLDVTDADSSRGGVSSLSGRGFGVGELRLQPADRQAMDSNSGLDVTDADSSRGGVSSLSGRGFGVGELRLVDSPCSIRLETLALNLRR

>Patinopecten_yessoensis_GnRH/CRZ (ALS92801.1)

MSSYTQILVAQLLLAGLLVAVVSGQNFHYSNGWQPGKRAPMMTSGTQLCSFRPHIKALLLWIIEDEVKRIKSCGSSGYDDIINLLQSKQSGLPLSSGMPSDSQ

>Bithynia_siamensis_goniomphalos_GnRH/CRZ (from EST, GAGS01040432.1)

MVRPVTTPSLSSFLLAACSCLMLMLLLLACPLPASAQSYHYSNGWNPGKRGIYGAASGPAVTDSGADDDLCLFRPKVLAFISKAIMEEISRIEQLCFPVDAVSGLKALLQQTAAAQSSSSAGPKLPEDGAKASW

>Crassostrea gigas_GnRH/CRZ (ADZ17180.1)

MKVSPCTQVIVMVLTLGLLCEVHAQNYHFSNGWQPGKRSYRGCTVRPEIRSILIKIIEDEVERIQKCSHSNIEDVFSLIQEKTGVDAREV

>Lottia_gigantea_GnRH/CRZ (from EST, FC805607.1)

MMPVPLKYFGLALTLALVTELAVGQHYHFSNGWKSGRKRSGGVSNLCEMRPELINYINTLLSEELNRIKNTCNLNTEDKDSDVDFSEGAFSRLQNGLKLAADRKWKK

>Octopus_vulgaris_GnRH/CRZ (BAB86782.1)

MSATASTTSSRKMAFFIFSMLLLSLCLQTQAQNYHFSNGWHPGGKRSALSDIQCHFRQQTKALIEKILDEEINRIITTCTGPVNEIADL

>Ruditapes_philippinarum_GnRH/CRZ (from EST, GAEH01003426.1)

MNACILLTTLVTMITIEKVQGQSYHFSNGWNPGKRSMQEPVCHFRQDVQTLVLKLIEDEVYRMLSDPSCIGGVPTLRNFLKKDLAYTVPLDDKK

>Capitella_teleta_GnRH/CRZ (from WGS, AMQN01007383.1)

MYRLELGCICSFCFCLLLCLACLQGCQGQAYHFSHGWFPGRKRSAPVSEKSSLLSEFPTSHGYAPIRSQKSSEIMDFFQYKRPIQRDDTPSTFDKEMCRMRPHVQQMIQDLINIEARRMAVECKSTIESAESSQQQREEDYPIKALEDWLRRGQGRLRRMESIRDESNE

>Danio_renio_GnRH (AAL99294)

MEWKGRLLVQLLLLVCVLEVSLCQHWSYGWLPGGKRSVGEMEATFRMLDPGDTVLSIPADSPMEQLSPIHIVNEVDAEGLPLKGQRYSDRRGRV

>Homo_sapiens_GnRH1 (NP_000816.4)

MCLRMKPIQKLLAGLILLTWCVEGCSSQHWSYGLRPGGKRDAENLIDSFQEIVKEVGQLAETQRFECTTHQPRSPLRDLKGALESLIEEETGQKKI

>Gallus_gallus_GnRH1 (NP_001074346.1)

MEKSRKILVGVLLFTASVAICLAQHWSYGLQPGGKRNAENLVESFQEIANEMESLGEGQKAECPGSYQHPRLSDLKETMASLIEGEARRKEI

>Gallus_gallus_GnRH2 (AEZ51861.1)

MAPGGCLLLALLLLAGTAQGQHWSHGWYPGGKRDLSAPQAIPAWSPWLVPSRGRKSRYDPPAKAKASRLKVPPPVDPAELYVVTERYRQHRLVLSALRSIFRSEVVQRKREAQRDAEDSAALSEEHRLLMAWNDAENARQRARR

>Cionia_intestinalis_GnRH6 (AAP06794)

MLDIEKDELAALLQRENSAFRDLLYHKNAGNFEKSDSGKFGSLKPQNNFPHLDLGLGVDLDAVDQWNRYKQANAQRMQDLGVPVNARQHWSYEFMPGGRRAAWENANVGVPVSRQHWSYEYMPGGRRSAGQHAMTKRQHWSKGYSPGGKKRSVDLSEFDDQGRRITKHEGMPEEPFKVEQPRPRNGIHGPAGLDQNEPDWKNWMNEQPAVSSDDKGTDVE

>Strangylocentrotus_purpuratus_GnRH (XP_800179.1)

MKQIITSLVSISAALLLFVLISEYTPRCNGQVHHRFSGWRPGGKKRSDAAEVNSNKITIERPQLPICQTTEERQLLEGDSDILGDLRRAANRMRLLQLFNLSKTRLNDLNDATSNEVDERPVYGDYLGTGL
